# Gene-metabolite annotation with shortest reactional distance enhances metabolite genome-wide association studies results

**DOI:** 10.1101/2023.03.22.533869

**Authors:** Cantin Baron, Sarah Cherkaoui, Sandra Therrien-Laperriere, Yann Ilboudo, Raphaël Poujol, Pamela Mehanna, Melanie E. Garrett, Marilyn J. Telen, Allison E. Ashley-Koch, Pablo Bartolucci, John D. Rioux, Guillaume Lettre, Christine Des Rosiers, Matthieu Ruiz, Julie G. Hussin

## Abstract

Studies combining metabolomics and genetics, known as metabolite genome-wide association studies (mGWAS), have provided valuable insights into our understanding of the genetic control of metabolite levels. However, the biological interpretation of these associations remains challenging due to a lack of existing tools to annotate mGWAS gene-metabolite pairs beyond the use of conservative statistical significance threshold. Here, we computed the shortest reactional distance (SRD) based on the curated knowledge of the KEGG database to explore its utility in enhancing the biological interpretation of results from three independent mGWAS, including a case study on sickle cell disease patients. Results show that, in reported mGWAS pairs, there is an excess of small SRD values and that SRD values and p-values significantly correlate, even beyond the standard conservative thresholds. The added-value of SRD annotation is shown for identification of potential false negative hits, exemplified by the finding of gene-metabolite associations with SRD ≤1 that did not reach standard genome-wide significance cut-off. The wider use of this statistic as an mGWAS annotation would prevent the exclusion of biologically relevant associations and can also identify errors or gaps in current metabolic pathway databases. Our findings highlight the SRD metric as an objective, quantitative and easy-to-compute annotation for gene-metabolite pairs that can be used to integrate statistical evidence to biological networks.

## INTRODUCTION

High throughput biotechnologies and analytic approaches applied to large human cohorts have recently revolutionized biomedical research, allowing the quantification and characterization of biological molecules to generate “omics” datasets. Genomics, the characterization of an individual’s DNA molecules, led to the identification of thousands of genetic variants associated with a trait, disease, or response to treatments, through large-scale genome-wide association studies (GWAS) [1]. Although GWAS have resulted in a better understanding of disease mechanisms, these approaches only consider genetic variation established at birth, and ignore the environment of an individual, influencing its biological state. An alternative strategy to complement traditional GWAS and better understand human biology is to comprehensively interrogate disease states at the molecular level using metabolomics, which offers a robust way to systematically measure thousands of low-molecular-weight compounds, called metabolites. After tissue extraction or collection of biological samples (usually blood and urine), metabolites can be detected, identified, and quantified using either mass spectrometry (MS) or nuclear magnetic resonance (NMR) [2]. As biomarkers of the underlying molecular dysfunctions, metabolite levels correspond to intermediate phenotypes (or endophenotypes) representing natural or clinical heterogeneity. Storing and sharing metabolomics knowledge is the focus of The Human Metabolome Database (HMDB), which is one of the largest and comprehensive curated collection of human metabolite and human metabolism data in the world [3].

Metabolomics can be seen as the study of the ultimate molecular response of an organism to genetic, environmental, and pathological modifications, but elucidating specific molecular mechanisms from metabolomics alone is difficult since many metabolites are involved in multiple biological processes. Linking metabolomics signals with genetics provides a promising approach to identify the implicated pathways and, as such, metabolite genome-wide association studies (mGWAS) are key approaches to integrate metabolomics with genomics. mGWAS report associations between metabolites and genetic loci, referred to as metabolite quantitative trait loci (mQTL), making it possible to study associations between millions of genetic variants and thousands of metabolites and to generate insightful hypotheses about uncharacterized regulatory mechanisms [2]. Although very powerful, interpretation of mGWAS results remains challenging. Indeed, millions of statistical tests are generally performed between single nucleotide polymorphisms (SNPs) and metabolites, leading to a huge multiple testing burden [4]. Furthermore, interpreting the biological meaning of an association between a given SNP and a metabolite requires interdisciplinary knowledge. In this context, the simplest way of prioritizing hypotheses remains to apply a conservative correction for multiple testing such as Bonferroni or Benjamini-Hochberg procedures [4]. The consequence of this correction is that only associations with the most significant p-values are reported whereas biologically relevant associations with a suggestive p-value may be missed.

With the increasing number of published mGWAS and their current limits, it is necessary to develop systematic methodologies to gain better insight into the mGWAS results by exploiting the most up-to-date biological knowledge. There are multiple resources available that aim at storing and describing known relationships between genes and phenotypes or endophenotypes (GWAS Catalog [5], PhenoScanner [6], OpenGWAS [7], Open Targets Genetics [8], PheLiGe [9], DisGeNET [10]) as well as tools to annotate gene-metabolites pairs based on the available resources but none of them have been specifically designed to address current mGWAS limitations [11, 12]. Only few methods have been developed to gain insight into the biological interpretation of mGWAS data specifically [13, 14], and none have been explicitly used to annotate mGWAS results based on well-curated biochemical knowledge, such as KEGG database [15]. KEGG is recognized for its high curation level of metabolic pathways for many model organisms, including humans, and is a reference for the reporting of enzymatic reactions, which is needed to understand the functional importance of gene-metabolite pairs. KEGG is a reference for the biochemical functions of genes with the descriptions of enzyme reactions, but a systematic and quantitative annotation procedure using this database in the context of mGWAS results remains to be explored. In this regard, recent studies have highlighted the applicability and utility of topological data analysis based on graph theory approaches [16], including the shortest path [17, 18], in revealing biological knowledge in the context of complex metabolic networks [13, 17, 18]. For example, the shortest path was used to characterize the impact of gene deletion on nearby metabolites within the metabolic network of E. coli [18]. It was also used to analyze the relationship between expression quantitative trait loci (eQTL) and metabolomic data, for example within the metabolic network of rat adipose tissues [17]. Still, this type of metric has not been applied to systematically annotate mGWAS results.

In this study, we assessed the utility of the shortest reactional distance (SRD) metric computed from KEGG database’s pathways to annotate mGWAS results and to help extract biological insights from them. We developed PathQuant, an R package, to enable a robust and systematic computation of SRD values between any lists of gene-metabolite pairs mapped onto KEGG graphs, which represent metabolic pathways, while keeping the original, well-curated, topology of each queried pathway. Focusing on genes encoding for enzymes and their associated metabolites, we applied the SRD annotation to two previously published mGWAS datasets in individuals from different ethnicities: an mGWAS [11] performed on 7,824 participants from the TwinsUK cohort [19] and the KORA study [20], referred herein as the TK study, and an mGWAS [21] performed on 614 Qatari participants from the Hamad Medical Corporation (HMC), referred herein as the HMC study. We explored results at varying levels of statistical significance. We found that the SRD metric enables identification of associations that do not meet currently accepted cut-off of statistical significance (suggestive associations) but which have high biological relevance. Finally, we performed an mGWAS on previously reported genetic and metabolomic data from a Sickle Cell Disease (SCD) cohort [22, 23], referred herein as the SCD study, and show how the SRD metric can be used to prioritize novel hits, including hits with a suggestive p-value.

## RESULTS

### Overview of study pipeline

PathQuant is a tool that converts a metabolic pathway map into a graph of biochemical reactions with metabolites as nodes and genes as edges, to compute the SRD path between a given gene-metabolite pair (STAR Methods, Supplementary Methods). To explore the potential of SRD values as an annotation metric to inform on the biological relevance of gene-metabolite pairs obtained from mGWAS, we developed a pipeline, presented in Figure 1. First, we gather mGWAS summary statistics and perform standard quality-control filters on genetic variants based on minor allele frequencies (MAF) and completeness. The second step of our pipeline only keeps bi-allelic SNPs at MAF > 0.01 (to exclude rare variants) and tested across all measured metabolites, but indels, tri-allelic SNPs and rare variants can be easily integrated, if present in mGWAS summary statistics. Third, SNPs are mapped to their closest genes coding for an enzyme (within 10,000 pb upstream and downstream) to obtain gene-metabolite pairs. As multiple SNPs will generally be linked with the same gene, only the minimum p-value of all SNPs is kept for a gene-metabolite pair, representing the strongest statistical signal for each pair within a study. These p-values can be categorized as either significant, suggestive, or non-significant according to appropriate thresholds that depend on the dataset (STAR Methods). We then retrieve KEGG IDs from all pairs (STAR Methods), and only retain pairs for which both IDs could be found. Third, we run PathQuant R package to compute the SRD value on all remaining pairs based on the KEGG overview graph (hsa01100). Note that other KEGG graph, or a combination of KEGG graphs, can be easily added at this step. SRD values obtained can be numerical when a path is found, can be given an infinite value (noted as “Inf”) when there is no path between the gene and metabolite in the queried graph, or an “NA” value when the gene or the metabolite (or both) are not found in the queried graph, despite having a valid KEGG ID. We can then perform several analyses of SRD values, as detailed below.

**Figure 1.**
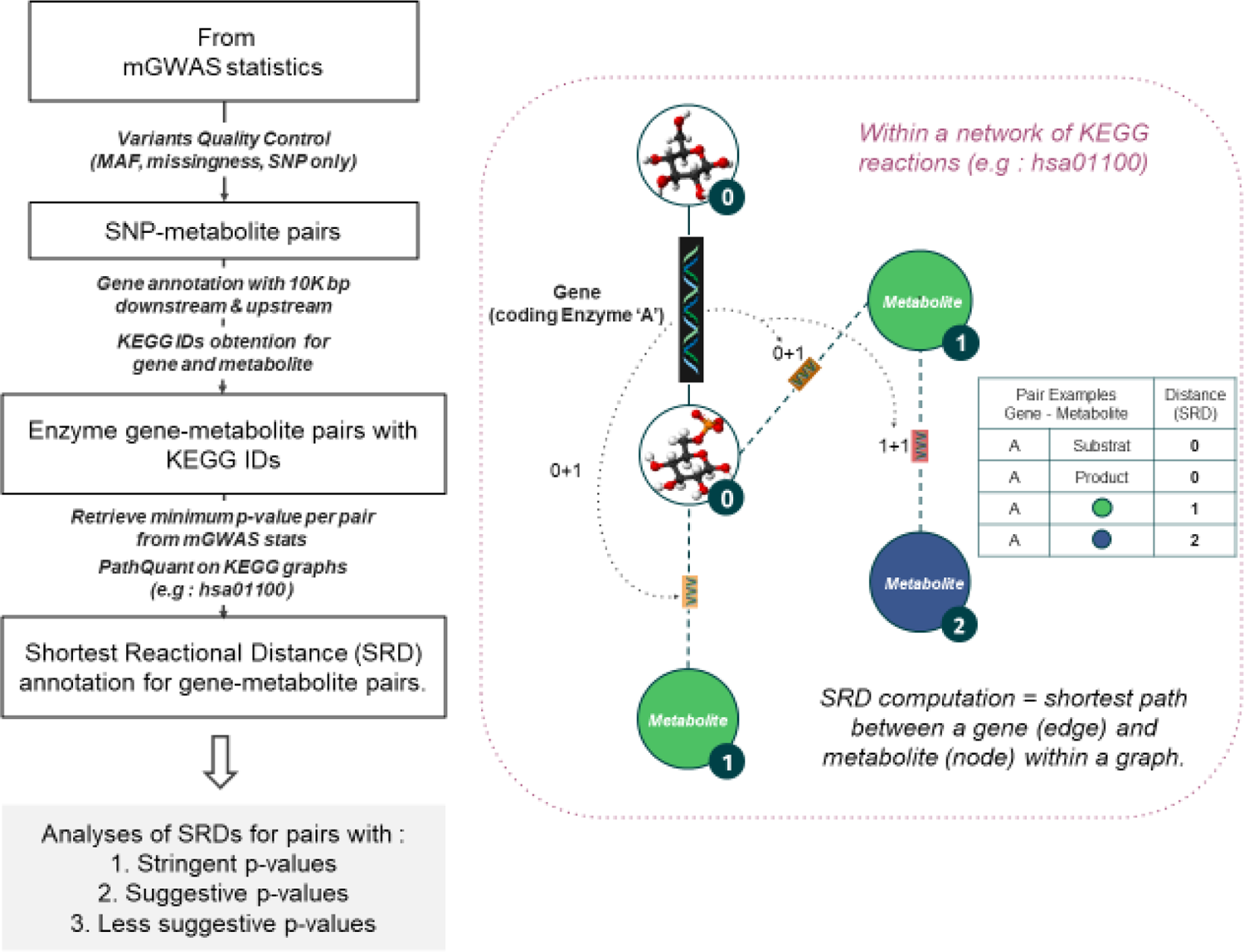
Overview of the Shortest Reactional Distance (SRD) annotation process. On the left panel, we show the main steps of the pipeline used to annotate mGWAS datasets. Boxes indicate input and outputs of processing steps described on arrows. The right panel shows the meaning of an SRD annotation of a reaction. The queried metabolite is in blue, the queried gene is Enzyme A and a sub-graph is shown. The SRD between Enzyme A and its main reactants, the substrate and the product, are of 0, everything running deeper adds 1 at each step (dashed arrows): green metabolites are at SRD = 1 and the queried metabolite is at SRD = 2 from Enzyme A.

### Stringent and suggestive associated pairs have shorter reactional distances

To test whether the SRD is a metric capable of capturing the biological relevance of a gene-metabolite pairs, we first explored the results of the TK study, as it represents one of the largest mGWAS studies conducted to date. This mGWAS was done on a group of participants from European descent, and many findings were replicated in follow-up studies, making it a well-validated dataset. Furthermore, it is considered a reference study by the community, resulting in an ideal dataset to test the hypothesis that stringently associated pairs will have lower SRD values. We started to explore this dataset by using the 74 gene-metabolite pairs with KEGG IDs passing the genome-wide significance cut-off set by the original authors [11] (Table 1). Among these, 40 gene-metabolite pairs were mapped onto KEGG overview graph. After excluding the two pairs with infinite SRD values, the median SRD value for the remaining 38 pairs is 1, which indicates a close biological relationship between the genes and their associated metabolites (Figure 2). To assess how significant this result is, we used two strategies: we compared the SRD values to (1) a null distribution (STAR Methods), built from all possible pairs of gene-metabolites that are present on the KEGG overview graph (Figure 2A); (2) to a permuted set of gene-metabolite pairs (Figure 2B), built with the 33 metabolites and 27 genes involved in the 40 significant gene-metabolite pairs (Figure 2C). There is a statistically significant difference between the TK pairs’ SRD values and the null distribution (Welch test, p-value < 3.48 × 10^−16^), confirming the close biological relationship between mGWAS gene-metabolite pairs in the TK study. Similarly, the median SRD value for the permuted pairs is 8, which is significantly higher than the median SRD of 1 (empirical p-value = 0.035). These results confirm that closely connected genes and metabolites are enriched in gene-metabolite pairs discovered in mGWAS.

**Figure 2.**
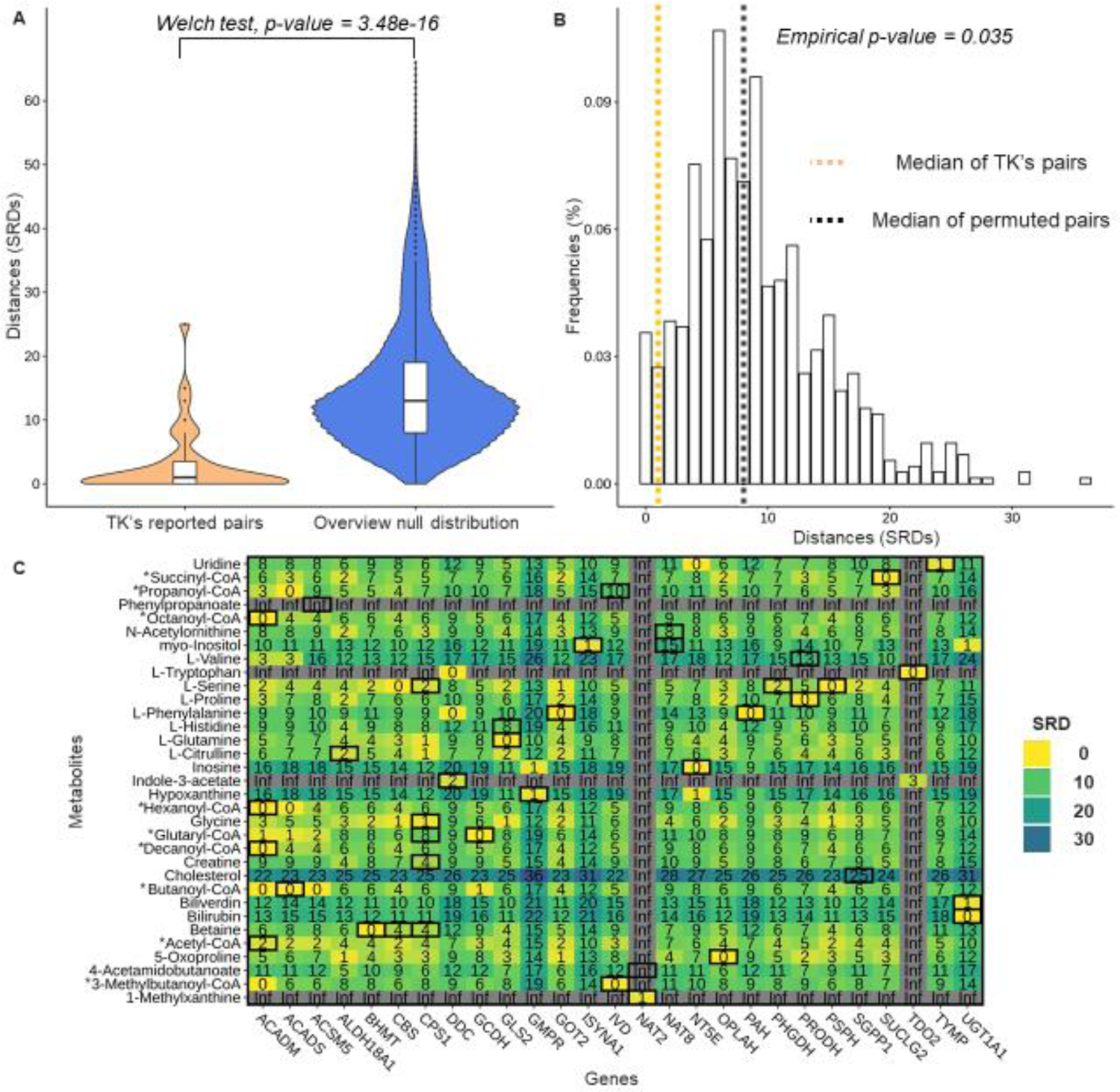
SRD of stringently associated gene-metabolite pairs in the TK study. (A) Comparison of SRD annotations for reported genome-wide significant associations from the TK study (orange) and the distribution of all SRD values within KEGG overview graph (hsa01100) (B) Distribution of SRD values computed from permuted gene-metabolite pairs from TK study, with a median SRD of 8 (black dotted line). The median SRD of 1 (orange dotted line) represents an empirical p-value of p=0.035. (C) Heatmap representing all genes and metabolites included in reported genome-wide significant associations from the TK study. The 40 mapped gene-metabolite associations are enclosed in black boxes on the heatmap (n=40). Abbreviations: CoA = Coenzyme A. * CoA metabolites are proxies for measured carnitines.

**Table 1.** Dataset description of used dataset for the TK, HMC, and SCD studies. Line 1: Dataset label for TK, HMC and SCD studies and the TK dataset without p-values cut-off (TK-None). Line 2: Maximum p-value cut-off. Line 3: Maximum number of samples available for the mGWAS study. Line 4: Number of SNP-metabolites pairs. Line 5: Number of gene coding for enzyme-metabolite pairs for which KEGG IDs have been obtained. Line 6: Number of SRD annotation within map hsa01100 (KEGG overview graph) categorized in three categories. Line 4,5,6 have the count of unique genes and metabolites involved in pairs between parenthesis. **\*:** the number of unique genes and metabolites are ignoring NA values. Abbreviations: ID = Identifier; SNP = Single-nucleotide polymorphism; SRD = Shortest Reactional Distance; Inf = Infinite value; NA = Not available, a missing annotation.

| Study name | TK | HMC | SCD | TK-None |
| --- | --- | --- | --- | --- |
| Summary stats cut-off | $1.03 \times 10^{-10}$ | $1.79 \times 10^{-7}$ | None | None |
| Sample size | 7,824 | 614 | 651 | 7,824 |
| SNP-Metabolite name pairs<br>(unique SNP - unique metabolites) | 336<br>(218-186) | 6 465<br>(3 192-100) | 3 139 070 208<br>(24 523 986-128) | 1 236 909 025<br>(2 617 408-486) |
| Enzyme Gene-Metabolite pairs with KEGG IDs: PathQuant input<br>(unique genes - unique metabolites) | 74<br>(43 - 56) | 68<br>(32-20) | 209 645 290<br>(3875-118) | 46 779 684<br>(3815-177) |
| Pairs' SRD annotations within KEGG Overview Graph<br>(unique genes - unique metabolites*) | SRD: 38<br>Inf: 2<br>NA: 34<br>(27 - 33) | SRD: 38<br>Inf: 4<br>NA: 26<br>(13 - 8) | SRD: 82 145<br>Inf: 38 980<br>NA: 336 125<br>(1275 - 95) | SRD: 86 219<br>Inf: 55 822<br>NA: 533 214<br>(1257 - 113) |

To visualize the relationship between all genes and metabolites involved in the significant pairs reported in the TK study within KEGG overview graph, we used a heatmap representation of SRD values (Figure 2C), highlighting the reported significant pairs using thick black boxes. Seventeen of these associations have an SRD of 0, implying that these gene-metabolite pairs are from a single enzymatic reaction. For example, PSPH (phosphoserine phosphatase) is catalysing the formation of L-Serine and results in an SRD of 0. Interestingly, within the 40 associations with an SRD annotation, some genes are closely connected to multiple metabolites, such as ACADM and CPS1 that have multiple small SRD values. Moreover, some genes are isolated and appear to be part of disconnected subgraphs of the KEGG overview graph, such as GMPR (Guanosine Monophosphate Reductase), NAT2 (N-Acetyltransferase 2) and TDO2 (Tryptophan 2,3-Dioxygenase), which obtained infinite SRD values with most metabolites.

The design of mGWAS can be heterogeneous, with distinct genomic and metabolomic technologies and pre-processing steps used across studies. To demonstrate the applicability of our annotation more broadly, it is important to use an independent study to replicate the observations above. Furthermore, there is a widely recognized bias towards white Europeans in genetics studies, which often makes results less generalizable in other ethnicities [24, 25], highlighting a need for new bioinformatics solutions to be tested in underrepresented populations. In line with these criteria, we further tested our approach in the HMC study, which has been performed in a different ethnic group, on Qatari individuals. In this mGWAS, the genomic data comes from the exome sequencing technology compared to genotyping data in the TK study, and the metabolomics data is generated using differing pre-processing steps (see STAR Methods). Furthermore, the authors of this study identified SNP-metabolite pairs with suggestive p-values, which were made available (Supplementary File number 7 of [21]), allowing us to explore whether including the suggestive mGWAS association results replicated the observation from the TK study. Of 68 gene-metabolite associations (Table 1) at a cut-off of p ≤ 1.4 × 10^−7^, 42 were mapped to the KEGG overview graph for which SRD values were computed. Four pairs had infinite SRD values (see Supplementary Figure 1C), while the 38 gene-metabolite remaining pairs (Table 1) had SRDs of 0 or 1 (Supplementary Figure 1), meaning that we have associations that are either substrate-product associations or with one intermediate step. Thus, we replicated the observation seen in the TK study of a significant enrichment of low SRD values compared to the null distribution in this second mGWAS (Welch test, p-value < 7.91 × 10^−60^, Supplementary Figure 1), even when considering a lower significant threshold than standard genome-wide cut-offs.

### SRD annotation can identify false negative hits

By using two different studies and different summary statistics cut-offs, we have determined that associations with stringent and suggestive p-values are enriched for low SRDs. We next explored more formally the relationship between the SRD values and the p-values from the TK study. We graphically defined the gene-metabolite significance cut-off at p < 3.16 × 10^−4^ of the TK study based on a QQplot (Supplementary Figure 2B, STAR Methods) and used it as a threshold for the minimum value to determine the relationship between SRDs and p-values. We observed a significant negative correlation between the mGWAS p-values and SRD values (R = −0.2, p = 9.1× 10^−11^, Figure 3A), meaning that the higher the significance of an association between a metabolite and a gene is, the closer they are likely to be in the KEGG overview graph. This negative correlation confirms the enrichment of biologically relevant pairs in significant associations, and suggests that combining p-values and SRD may help prioritizing associations of high biological interest. Furthermore, the SRD value could be used to select for further analysis pairs with small SRD having low p-value that nevertheless do not pass stringent significance cut-offs.

**Figure 3.**
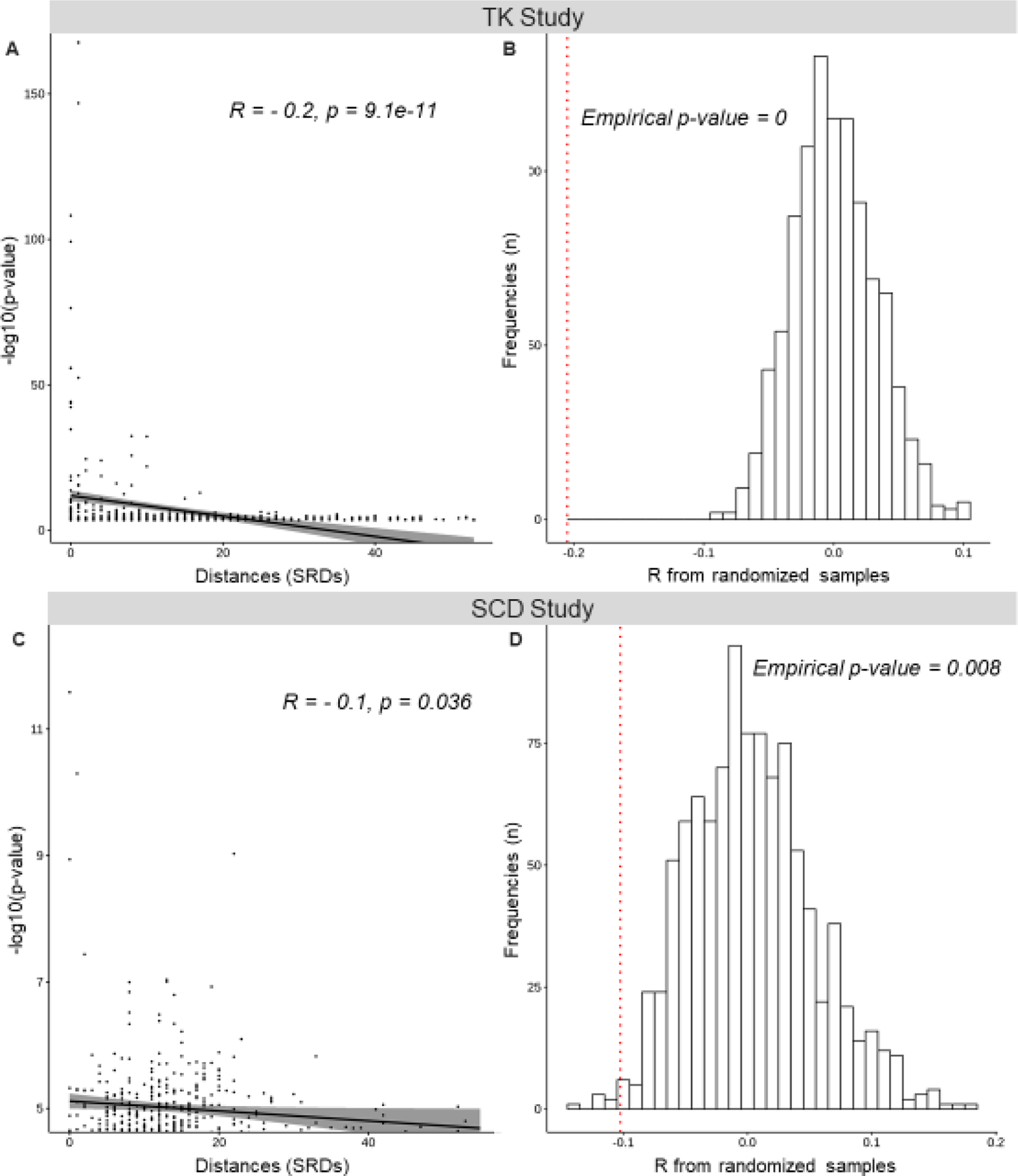
Relationship between p-values and SRD values for TK and SCD studies. Correlation plot between the –log10(p-values) and SRD values for gene-metabolite pairs in (A) the TK study (p-value cutoff was 3.16 × 10^-4^, Supplementary Figure 3B) and (C) the SCD study (p-value cutoff was 3.16 × 10^-5^, Supplementary Figure 3E). Correlation computed on permuted data (N=1000) to take into account the graph structure allowed computation of empirical p-values for TK (B) and SCD (C), with the true correlation coefficient presented (dotted red line).

As a result of multiple testing burden between each variant and each metabolite level, only top hits are generally reported in mGWAS, but it is well known that there could be false negative hits [26]. Thereby, we tested the potential for the SRD metric to identify some of these false-negative candidates in these published mGWAS. In the TK study, we focused on the genes and metabolites involved in stringent associations (presented in the heatmap, Figure 2C), which include 27 genes and 33 metabolites. We noticed gene-metabolite pairs annotated with a smaller SRD than the initially reported association, such as the ALDH18A1 (aldehyde dehydrogenase 18 family member A1) and L-citrulline pair, which has an SRD value of 2, whereas other pairs involving L-citrulline show smaller SRD values. For instance, the CPS1 (carbamoyl-phosphate synthase 1) and L-citrulline pair has an SRD = 1 but did not pass the stringent p-value cut-offs of the TK study, despite a p-value of 1.749 × 10^−8^. The SRD value of 1 reflects a sequence of reactions pertaining to the urea cycle catalysed by CPS1 for the formation carbamoyl phosphate from ammonia and bicarbonate and by the ornithine transcarbamylase (OTC) for the formation of L-citrulline from carbamoyl phosphate and L-ornithine [27]. The association between CPS1 and L-citrulline has recently been described twice in the literature with a p-value of 1 × 10^−25^ with SNP rs1509820 [28] and 1 × 10^−14^ with SNP rs975530777 [29]. The fact that this association was not significant in the TK study can thus be considered a false negative result.

Using the same approach, we also identified an example within the HMC study of a gene-metabolite pair at SRD = 0, which was not reported as a significant association because of a suggestive p-value (p =1.17 × 10^-7^). This association is between UMPS (uridine monophosphate synthetase) and orotate (Supplementary Figure 1C, indicated in red). UMPS is a bifunctional enzyme that is part of the *de novo* pyrimidine biosynthetic pathway; its orotate phosphoribosyltransferase subunit catalyses the addition of ribose-5-phosphate to orotate to form orotidine monophosphate (KEGG reaction ID R01870). Given that a significant association has been previously reported between UMPS gene and orotate [30], we could consider that this finding is replicated in the HMC study but only by adding information about its SRD value.

Taken altogether, the examples highlighted the opportunity to explore associations annotated with low SRD values in mGWAS, beyond those with stringent significance cut-offs. Identifying these potential false negative hits illustrates how useful the SRD annotation can be in discovering biologically relevant results which would be missed when only considering stringent p-value cut-offs in mGWAS reporting.

### Case study: mGWAS in Sickle Cell Disease patients

To exemplify how the SRD metric can be used to prioritize mGWAS results, we present a new mGWAS analysis performed in sickle cell disease (SCD) patients of African or African-American ancestry. While the cause of SCD has been known for over a century, the molecular determinants of the severity of this blood disease remain unknown and are influenced by genetic variants unlinked to the beta-globin gene [31]. An mGWAS was performed in 651 SCD patients, quantifying a total of 128 metabolites, to identify metabolites associated with genetic markers. A total of 165 unique SNPs (with MAF > 1%) passed a genome-wide significance cut-off of p < 7.8125 × 10^−10^ (Supplementary Figure 2, 1 × 10^−7^/ 128 metabolites) and an additional unique 256 SNPs passed a suggestive cut-off of p < 1 × 10^−7^ (Supplementary Figure 2C). We identified a total of eight loci associated with metabolite levels (Supplementary Figure 3), with four of them reported in previous studies (Supplementary Table 2). The four additional associations were not previously reported neither in the TK/HMC studies, nor in a large mGWAS of human blood metabolites [32]. In all four cases, the frequency of the top associated SNP is larger in individuals of African descent than in Europeans according to the gnomAD Genomes database, but these hits were not found in the three largest mGWAS done in individuals of African ancestry to date [33–35]. We recognize that while interesting, these novel associations will need replication in an independent cohort.

We generated a QQplot based on minimum p-values for the gene-metabolite pairs extracted from the SCD mGWAS (STAR Methods) and defined the gene-metabolite significance cut-off at p < 3.16 × 10^−5^ (Supplementary Figure 2D). We observed a significant negative correlation between the mGWAS p-values and SRD values (R = −0.1, p = 0.036), replicating the result observed in the TK study, demonstrating that this relationship between significance and SRD is reproducible in a disease cohort, where biological mechanisms can be altered (Figure 3C). Given these observations, we investigated whether associations above the gene-metabolite significance cut-off of p < 3.16 × 10^−5^, but below genome-wide significant cut-offs usually required to report associations, could be prioritized according to SRD values (candidate pairs below the plain line in Figure 4). We split the associations into two suggestively significant categories, one that includes association p-values between the genome-wide significance cut-off and p < 1 × 10^−7^ (Supplementary Figure 2C) and another that includes associations p-values between p < 1 × 10^−7^ and the gene-metabolite significance cut-off of p < 3.16 × 10^−5^ (Supplementary Figure 2D), which we labelled S+ and S-, respectively. To evaluate the potential of association results in each significance categories, we aimed to compute the proportion of small SRD values. From the distribution of all SRD values computed on the KEGG overview graph (Methods), we observed that 25% of SRD values (first quartile) are lower or equal to 8 (Supplementary Figure 4). We thus used the SRD value ≤ 8 as a threshold for considering SRD values as small, reflecting close biological relationships based on the graph topology. Furthermore, 90% of gene-metabolite significant pairs in the well-powered TK and HMC studies have SRD ≤ 8 and, out of the genome-wide significant associations with a numerical SRD value in this mGWAS (Block R, Figure 4), two out of three have SRD ≤ 8.

**Figure 4.**
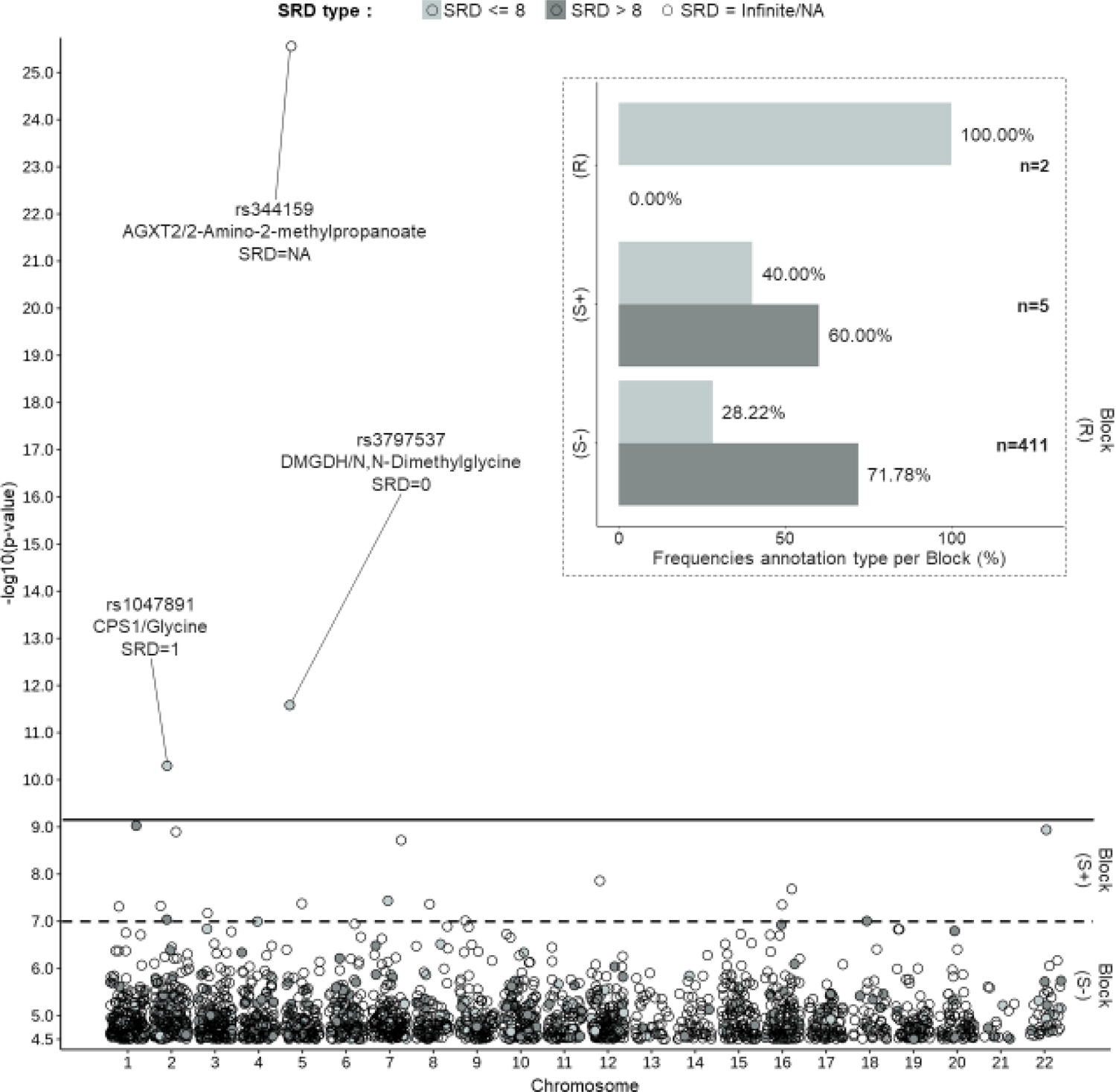
Identification of potential false negative hits for different p-values cut-offs in the SCD study. The main panel represents -log10(p-values) for gene-metabolite pairs for SNPs on each chromosome (x-axis). Coloring represents the SRD annotation from 3 categories: SRD = Infinite or NA (white), numerical SRD ≤ 8 (light grey), numerical SRD > 8 (dark grey). On the top right panel, the histogram represents the frequencies of SRD ≤ 8 and > 8 for each block. The frequency percentage is obtained by summing only the pairs with a numerical SRD value (n) within each category according to different p-value cut-offs: R: p < 7.8125 × 10^−10^, S+: 7.8125 × 10^−10^ < p < 1 × 10^−7^ and S-: 1 × 10^−7^ < p < 3.16 × 10^−5^.

Two (40%) out of five associations from the S+ category have small SRD values, and 116/411 for the S-category (28.22%). Next, we manually investigated the 118 suggestive associations with SRD ≤ 8 by searching the literature for these gene-metabolite pairs and found six previously described associations (5%): PRODH (proline dehydrogenase 1) and L-proline [11, 36, 37], NT5C3A (5’-nucleotidase, cytosolic IIIA) and orotate [30], GATM (glycine amidinotransferase) and creatine [38], UPB1 (beta-ureidopropionase 1) and 3-ureidopropionate [32], UGT2B17 (UDP glucuronosyltransferase family 2 member B17) and sn-glycerol 3-phosphate (previously described for total phosphoglycerides) [39], DGKH (diacylglycerol kinase eta) and tetradecanoyl-carnitine [40], implying that these are likely to be real associations in SCD patients (Table 2). We thus estimate that at least 5% of associations in this category are false negative and that the SRD metric has added value in identifying them. It also demonstrates the ability of the SRD metric to retain pairs that would not be otherwise reported, thus improving the mGWAS’ potential for discovery.

**Table 2:**
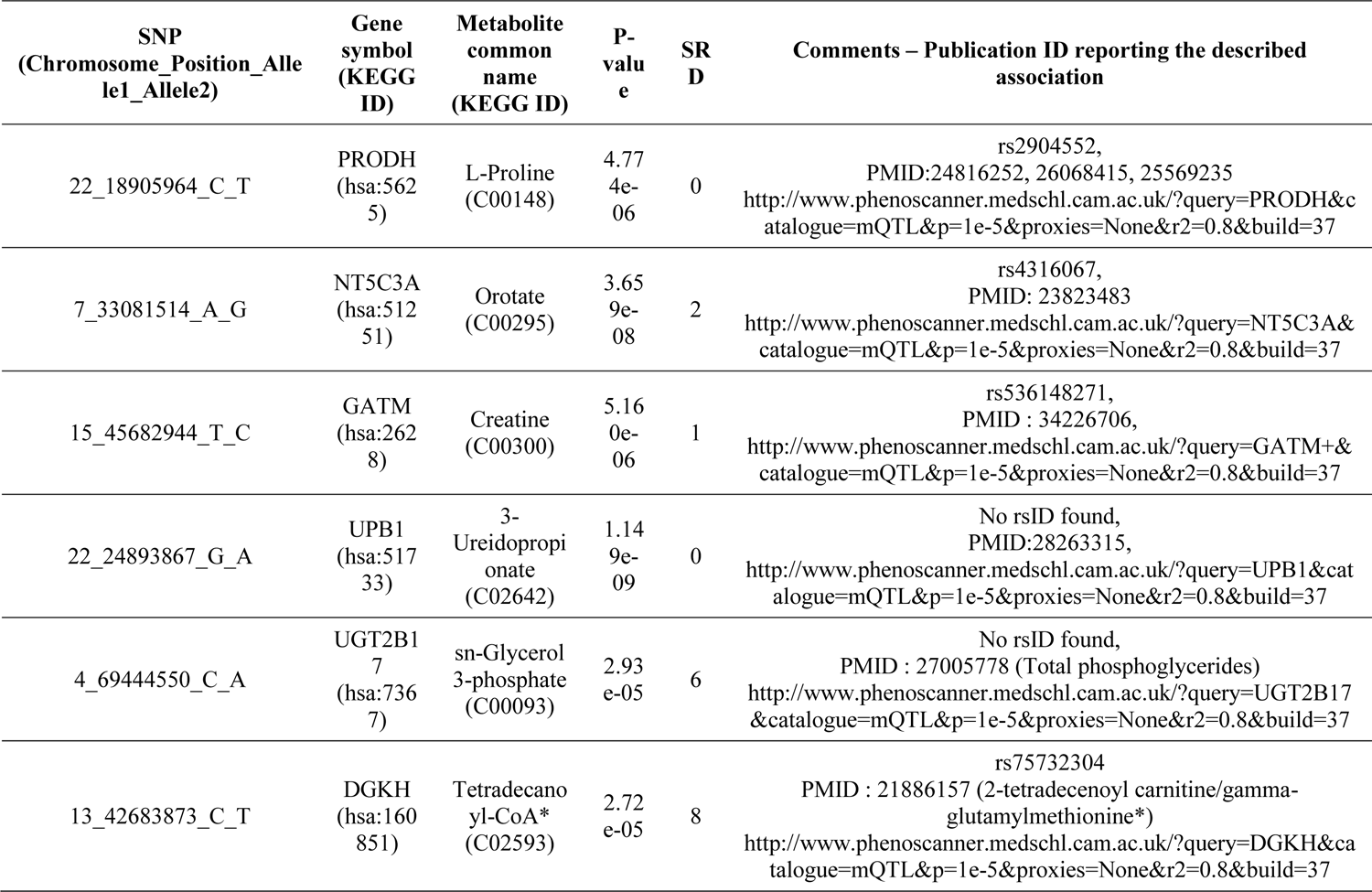
Associations with SRD lower or equal to 8 for the SCD study, illustrating potential false negative hits discovered by manual verification. Column 1: SNP information formatted as CHROMOSOME_POSITION_ALLELE1_ALLELE2 in hg19. Column 2: Gene symbol found the in the KEGG ID entry. Column 3: Metabolite common name found the in the KEGG ID entry. Column 4: P-value of the association between the SNP and the metabolite. Column 5: SRD annotation of the gene-metabolite pair using KEGG graph hsa01100. Column 6: Miscellaneous information for the association: rsID if available; PMID of the publications reporting association between the gene, or the SNP with the associated metabolite; link to the query used in Phenoscanner website. Abbreviations: ID = Identifier; SNP = Single-nucleotide polymorphism; SRD = Shortest Reactional Distance, PMID = PubMed Identifier. * CoA are representatives of the measured carnitines.

## DISCUSSION

The identification of an association between a SNP and a metabolite is usually supported solely by the p-values of the statistical test without consideration for the known metabolic pathways. Although the possibility of false positive hits is well understood among geneticists following up on mGWAS results, the fact that reporting of gene-metabolite pairs using only statistical significance can lead to an incomplete list of associations is less discussed. In this study, we demonstrate how a simple metric called shortest reactional distance (SRD) can be useful for the reporting and the *ad-hoc* annotation of gene-metabolite pairs for several p-values acceptance cut-offs. By using previously published mGWAS, we have shown that the SRD is a metric capable of retrospectively identifying a number of false negatives. The datasets we used in our study involved different genomics and metabolomics protocols, different preprocessing strategies, different sets of genetic variants (genotyped, imputed, exclusive to exons) and of metabolites, different ethnicities (from European, African and Middle Eastern ancestries), disease-based and population cohorts, illustrating the wide range of contexts in which our annotation is applicable.

PathQuant, the package developed to compute the SRD metric in this study, is not the only package available that computes the shortest path metric between genes and metabolites. Similar to PathQuant, MetaboSignal [13] is an R package based on the same mathematical criterion of shortest path metric and also uses metabolic pathways from the KEGG database. There are, however, important differences between these two methods. MetaboSignal combines both metabolic and signaling pathway maps. While this can provide novel information about the interaction between genes and metabolites that goes beyond the known enzymatic reactions and pathways, it modifies the original topology of the curated pathways (signaling and metabolic). These changes are not necessarily supported by biological data, which is crucial to ensure their validity, as emphasized by Dumas et al. [17]. Thus, for our assessment of the potential of SRD annotation in mGWAS, we favored a simpler approach that focuses only on curated metabolic pathways, by converting a metabolic pathway map from KEGG into a graph of biochemical reactions with metabolites as nodes and genes as edges. PathQuant computes SRD values from any given list of gene-metabolite pairs using any given metabolic pathway graph in KGML format thus keeping the original and curated topology of the metabolic pathways reported by KEGG. Of note, a direct comparison of the shortest path metrics calculated by PathQuant and MetaboSignal on the list of gene-metabolite pairs of the mGWAS investigated here could not be performed, as the KEGG overview pathway graph we used with PathQuant is not accepted as input by MetaboSignal. Indeed, MetaboSignal computes its shortest path metric using a custom graph that is built from a pre-selected list of signaling and metabolic pathways.

The computation of the SRD metric with PathQuant within KEGG graphs leads to several possible values: numerical, infinite or NA. Having a numerical annotation to a pair which is based on known pathways leads to a better understanding of the underlying biology of an association. Indeed, we have shown that SRD values decrease as statistical significance of gene-metabolite pairs increases: this negative correlation between the level of significance and SRD values suggests an enrichment of biologically relevant pairs as the p-value decreases, thus making SRD a promising annotation metric to improve mGWAS reporting. We defined two categories for numerical values (short SRD: 0 < SRD ≤ 8; large SRD: SRD > 8), which have been derived from the topological properties of the KEGG overview graph (ID: hsa01100) as this graph contains all curated human biological reactions, but different thresholds could be used in different contexts. The annotation of a pair with an SRD lower or equal to 8 suggests that the gene, its expression levels or the protein it codes for, may have a direct influence on the associated metabolite concentration. This is illustrated by the statistical difference between the SRD values of mGWAS pairs and the null distribution of SRD values within the KEGG overview graph and by manual investigation of specific pairs of interest, such as the CPS1 and L-citrulline finding (SRD=1) in TK study, the UMPS and orotate example (SRD=0) in HMC study. Moreover, for the SCD study, the gene-metabolite pairs presented in Table 2 are annotated with an SRD lower or equal than 8 and have been already reported to be associated with the corresponding metabolite levels in the literature. Despite the number of investigated associations being very small for the R and S+ categories, we observed a decreasing proportion of pairs with values lower than 8 and an increasing frequency of pairs with values greater than 8, as p-values increases, in line with the negative correlation observed. These results demonstrate, across all different mGWAS datasets we used, the potential of using the SRD metric with a cut-off of SRD ≤ 8 in order to identify false negatives hits and prioritize follow up studies and experiments.

Another interesting case is when the SRD value is large, but the p-value is highly significant. In this case, a direct influence of the gene on the associated metabolite is less clear. For example, we noticed a large SRD value for the association between the alkaline phosphate (ALPL) and the phosphocreatine (SRD = 22; p-value = 9.356× 10^−10^). One possibility is that this association could have been a false negative. However, digging further into the literature, we found that phosphocreatine can, in fact, be the substrate of the enzyme encoded by the ALPL gene [41, 42]. Thus, the true SRD value for ALPL-phosphocreatine pair should be 0, and the high SRD value computed is a consequence of the current state of the publicly shared knowledge available on KEGG. This result highlights a limitation of our approach, which is the incompleteness of some metabolic pathways in KEGG. However, our SRD annotation pipeline of mGWAS results can help identify those cases and could be useful to detect gaps and errors in metabolic network databases. Other cases of SRD > 8 for highly significant association have been noted in our work, notably in TK study (Figure 2A, Figure 2C; Figure 3A). When the path computed in KEGG is accurate and the association is independently replicated, higher values of SRD can identify metabolic pathways involving more indirect gene-metabolite relationships. By themselves, these cases are of interest as they illustrate a violation of the hypothesis that biological proximity between a gene and a metabolite is needed to regulate its level.

An SRD annotation with an infinite or NA value means that there was no path between the gene and the metabolite within the chosen graph, because they are not connected (infinite value) or because one of the two entities (or both) are missing from the queried graph (NA value). In most cases, there likely exist no biological paths between these entities, but if the association is highly significant, it could reflect actual gaps within the graph again. Investigation of enzymes or metabolites with an unusual number of infinite or NA values may lead to more complete metabolic pathway databases. Beyond these potential gaps in KEGG, other limitations of this resource are the metabolic reactions provided do not specify cofactors, enzymatic complexes required for the reaction to happen, and directionality, resulting in a decreasing level of complexity and accuracy of the metabolic pathways. Thus, computing SRD on other resources for pathway mapping, such as Recon3D, could improve our distance-based annotation pipeline. Indeed, this resource is considered the most complete reconstruction of human metabolism so far [43, 44]. Furthermore, in contrast to KEGG, Recon3D has the advantage of including more genes encoding transporters, which would allow to increase the breath of SRD computation beyond enzymatic reactions. It also includes information about cellular compartments and the directionality of reactions is known, in contrast to KEGG graphs. Despite these advantages, the available file format (sbml) used in Recon3D to represent metabolic pathways is incompatible with the graph structure we used for KEGG. Given that the goal of this study was to establish if the SRD annotation is useful for mGWAS annotation, a simple representation of metabolic pathways is highly appropriate and it is outside the scope of the present study to implement an SRD metric based on Recon3D. Future implementations on alternative databases should consider two additional desirable characteristics of KEGG when comparing results: first, KEGG is the only database for which pathway maps were built on known reactions from humans and others species, a clear advantage for non-human mGWAS studies [45, 46]; second, KEGG pathway maps are built with an explicit labeling of side compounds definition (such as ATP or H_2_O) from KEGG RPAIRS [47], making their removal from curated reactions possible, which eliminates shortcuts created by their over-representation within the graph when computing SRD [48].

A crucial step for the annotation of gene-metabolite pairs with SRD values is to obtain the KEGG IDs for the genes and metabolites. Although gene names and IDs are highly standardized [49] there is a lack of uniformed nomenclature for metabolites. In most cases, there are multiple ways to refer to a metabolite, with different studies using different reporting conventions, which complicated the re-analysis of published datasets. For HMC and SCD studies, HMDB IDs [3] were provided by the authors, which could be easily converted to KEGG IDs using the MetaboAnalyst Convert tool. In the case of the TK study, we performed a manual annotation of the common names to KEGG IDs, but this process was time consuming and required having the relevant expertise. This manual work allowed us to include listed metabolites that did not have KEGG IDs, such as acylcarnitines (see Material and Methods). These metabolites are measured in plasma but arise from intracellular metabolism of their acyl-CoA counterparts, which in contrast to acylcarnitines do not cross the plasma membrane. Based on the known direct link between the acylcarnitines and the acyl-CoAs we have used the CoA counterparts to represent these metabolites, which do have KEGG IDs. The increasing quantity of released studies involving metabolomics in the literature encourages the community to release new standards to refer metabolites, such as Simplified Molecular-Input LineEntry System [50], COordination of Standards in MetabOlomicS [51], IUPAC International Chemical Identifier [52]. These efforts for standardisation of metabolites naming, and the reporting improvements they allow, are leading to a promising leap for the field, as this will result into better links being made between novel findings and already published data.

In practice, SRD annotations can be a great addition to bioinformatics pipelines in order to reduce the number of potential candidate pairs to follow up on, by prioritizing hits based on a curated biological information, as the number of released mGWAS grows. Furthermore, this metric could be added within already available user-friendly resources such as mGWAS Explorer [14]. SRD values could also play a role in designing targeted studies, in a context where extensive mGWAS cannot be done or is not relevant, for example if the research question is about finding the genes or proteins that regulate the levels of a specific metabolite, or conversely, finding which metabolites are regulated by a specific enzyme. In this latter case, it would make sense to focus more specifically on the metabolites showing low SRD values with the targeted enzyme. In the future, we believe that extending the implementation by using other appropriate resources for pathway mapping would likely improve coverage and enhance the value of the SRD metric in mGWAS annotation pipelines.

In conclusion, we consider this work as a proof of concept of the benefits of using shortest reactional distance metrics for annotating mGWAS results based on a model representation of the human metabolism provided by the KEGG database. These metrics can also be used as a tool for metabolic databases in order to more easily identify gaps within current metabolic pathway graphs by using a new information provided by mGWAS. In this multi-omics era, we anticipate that large scale studies looking for associations between genomics, proteomics and metabolomics signals will soon become a new standard, as illustrated by recent studies in mice and humans [53, 54], resulting in additional protein-metabolite pairs, for which the computation of SRD values may add great value.

## STAR METHODS RESOURCE AVAILABILITY

### Lead contact

Further information and requests should be directed to and will be fulfilled by the lead contact, Julie Hussin.

### Materials availability

This study did not generate new materials.

### Data and code availability

- Source data statement. This paper analyzed existing, publicly available summary mGWAS data for the TK and HMC studies. The SCD study data comes from genomic and metabolomics of two previously published studies. Raw data can be made available upon request to GL. Summary statistics of SCD study will be made available publicly upon journal publication and from the lead contact upon request in the meantime.
- All additional files necessary to reproduce the reported results of this paper are available at: www.github.com/HussinLab/PathQuant/Publication/References.
- PathQuant source code is available at www.github.com/HussinLab/PathQuant/.
- Any additional information required to reanalyze the data reported in this paper is available from the lead contact upon request.

## METHOD DETAILS

### Overview of the mGWAS Datasets

In this study we used three different mGWAS datasets. The TK study refers to an mGWAS [11] previously performed on 7,824 participants (plasma or serum samples) from the TwinsUK cohort [19] and the KORA study [20]. The mGWAS was carried out on 2.1 million genotyped SNPs and 529 metabolites (assessed by targeted and untargeted metabolomics). The authors reported 299 SNP-metabolite (or ratio of metabolites) significant associations (at cut-off of p ≤ 1.03 × 10^−10^ for metabolite concentrations and p ≤ 5.08×10^−13^ for pairwise metabolite ratios) involving 187 unique metabolites and 145 loci, annotated as 132 causal genes [11]. The TK study also made available mGWAS output files from METAL software [55] involving 486 metabolites without any p-value cut-off [www.metabolomics.helmholtz-muenchen.de].

The HMC study refers to an mGWAS [21] previously performed on 614 Qatari participants (plasma samples) from the Hamad Medical Corporation. The mGWAS was carried out on 1.6 million imputed exome variants and 826 metabolites (assessed by targeted and untargeted metabolomics) and reported 3,127 significant associations (at cut-off of p ≤ 2.2 × 10^−10^ for both metabolite concentrations and pairwise metabolite ratios) in 21 locus-metabolite pairs. The suggestive association results were also provided to the community, reporting all associations with cut-off of p ≤ 1.4 × 10^−7^ (Supplementary Data 7 in [21]), which include 6517 SNP-metabolite (or ratio of metabolites) pairs with available p-values.

In the SCD study, an mGWAS was performed here using genetics and metabolomics data published in different studies [22, 23] on a total of 651 SCD patients (plasma samples) including 401 individuals of African ancestry in the Genetic Modifier (GEN-MOD) cohort and 250 African-American individuals from Southwest USA in the Duke University Outcome Modifying Genes (OMG) cohort. Metabolomics profiling was performed at the Broad Institute, and appropriate statistical modelling was used to account for residual batch effects [56]. Briefly, for association testing, 128 metabolites were profiled using a targeted approach in 651 plasma samples from SCD patients. Metabolites were inverse normal transformed, adjusting for age and sex. We then generated summary statistics for each cohort individually in a linear regression model, accounting for relatedness using a kinship matrix as implemented in rvtest (v. 20171009) [57] in GEN-MOD. The software SNPTEST [58] was employed in OMG to generate cohort-specific summary statistics. We then meta-analyzed the effect size estimates and standard errors from GEN-MOD and OMG using METAL [55]. After the pre-processing step, 6 million SNPs were kept.

### Mapping shortest reactional distances (SRD) onto KEGG

To compute the SRD metric, the R package PathQuant was developed [available at: www.github.com/HussinLab/PathQuant]. PathQuant converts a metabolic pathway map into a graph of biochemical reactions with metabolites as nodes and genes as edges (Figure 1). Briefly, PathQuant takes as input a list of gene-metabolite pairs as pairs of KEGG identifiers (IDs) and a list of metabolic pathways (eg. “hsa01100”, referred herein as KEGG overview graph, a global concatenation of multiple distinct pathways). PathQuant then uses a KEGG XML file format (KEGG Markup Language, KGML), downloaded using the KEGG API [www.kegg.jp/kegg/rest/keggapi.html] to build the most up-to-date KEGG undirected graphs. Next, PathQuant computes the SRD path between a gene and a metabolite from a given pair. The SRD values are obtained using the breadth-first search Dijkstra algorithm [59]. PathQuant outputs a text file containing genes and metabolites classification, Enzyme Commission number (EC), KEGG Brite IDs, KEGG IDs of metabolic pathways for the SRD computation, and the SRD values for all pairs. PathQuant also allows the visualization of SRD values annotation in a heatmap, leading to a better identification of potentially interesting hits.

### Extracting and annotating gene-metabolite pairs from mGWAS summary statistics

Using *bedtools version v2.30.0* [60], we annotated each SNP to genes with KEGG IDs, using a custom bed file: we downloaded gene coordinates for human genome build GRCh37.p13 from NCBI (NCBI Homo sapiens Updated Annotation Release 105.2020/10/22, gff format) and modified it to add 10,000 base pairs upstream and downstream for each of the 26,105 genes with KEGG IDs. Because we focused on pairs involving genes encoding enzymes, which are the most represented genes in the KEGG database, we only annotated SNPs with KEGG IDs classified as enzymes within the KEGG Brite database by using the Brite enzyme code “BR:hsa01000”, leading to a subset of 4,049/26,105 genes. For metabolites, if available, we used the provided IDs, either KEGG IDs directly or HMDB IDs. For metabolites with HMDB ID the list was queried to *Metaboanalyst* [61][www.metaboanalyst.ca/faces/upload/ConvertView.xhtml] to get the corresponding KEGG IDs. For metabolites without KEGG or HMDB IDs provided, KEGG IDs retrieval was achieved automatically by parsing the common names into the most standard KEGG name format, and then queried the metabolite names with new format to MetaboAnalyst. Additionally, specific metabolites were manually treated: acylcarnitines without KEGG IDs were swapped to their acyl-CoA counterparts when available in KEGG. For each gene-metabolite pair within a dataset (TK, HMC and SCD study), only one mGWAS p-value was kept, which is the minimum p-value obtained for the association between any SNP annotated to the gene and that specific metabolite. Based on that p-value, each pair is annotated as genome-wide significant, suggestive, or non-significant: genome-wide p-value cut-offs were derived from a standard GWAS Bonferroni correction approach (dividing by the number of tested metabolites). Suggestive significance cut-offs, were obtained graphically from quantile-quantile plots (QQplots) of minimum p-values for the variants-metabolite pairs of corresponding study. Specifically, we determined the value on the QQplot x-axis where the slope starts to increase drastically.

### SRD null distribution and investigation of candidate gene-metabolite pairs

To obtain a null distribution of SRD values from the KEGG overview graph (hsa01100), we gathered all different KEGG IDs for both metabolites and genes found within the graph, and then ran PathQuant on all possible pairs (Supplementary File 1). The KEGG overview graph version (KGML v0.7.2 file) includes 1,351 genes and 2,889 metabolites, generating a total of 3,903,039 pairs for which we computed the SRD values, referred herein as the KEGG overview graph’s SRD null distribution. The first quartile of this distribution is used as the threshold to categorize any pair with a close or far biological relationship label. This threshold was determined because 25% of the smallest values are below half of the mean, leaving 75% as a representative sample of distant relationships between genes and metabolites. We also performed a manual investigation of candidate gene-metabolite pairs (Table 1, Supplementary Table 1) by extracting the corresponding rsID using dbSNP annotation of UCSC Genome Browser [62], searching for published associations of the metabolite with (1) the SNP rsID within GWAS Catalogue and/or with (2) the gene symbol within PhenoScanner [6] (with parameters: cut-off p-value=1e-5, cut-off r^2^=0.8, build=37).

## Supporting information

Key Resources Table

## Acknowledgements

We thank all SCD participants for their contribution to this project. We thank members of the Hussin group for helpful discussions and advice throughout this project, specifically Jean-Christophe Grenier. We thank Fabien Jourdan, Clément Frainy and the iGenoMed Consortium for initial discussions on this project. This work was completed thanks to computational resources provided by Calcul Quebec clusters Narval, Beluga and Cedar, maintained by the Digital Research Alliance of Canada. CB was supported by a PhD Scholarship from the NSERC-CREATE Metabolomics Advanced Training and International Exchange (MATRIX) program. MR is a FRQS Junior 1 research scholar, JGH is FRQS Junior 2 research scholar, GL holds the Canada Research Chair (tier 1) in the genetics of heart and blood diseases, JDR holds a Canada Research Chair in Genetics and Genomic Medicine. This study was supported by funding from the Montreal Heart Institute Foundation and IVADO PRF Grant (PRF-2019-3378524797) to JGH and MR, and a National Sciences and Engineering Research Council (NSERC) Discovery Grant (RGPIN-2022-04262) to JGH, financial support of Génome Québec, Genome Canada, the government of Canada, and the *Ministère de l’enseignement supérieur, de la recherche, de la science et de la technologie du Québec*, the Canadian Institutes of Health Research (with contributions from the Institute of Infection and Immunity, the Institute of Genetics, and the Institute of Nutrition, Metabolism and Diabetes), Genome BC and Crohn’s Colitis Canada (via Genome Canada grant number GPH-129341) to CDR and JDR, Agilent Technologies Research Grant (3893 & 4075) to CDR. The work in the GEN-MOD cohort is funded by the Canadian Institutes of Health Research (CIHR, PJT #156248) and the Doris Duke Charitable Foundation. GEN-MOD sample and data collection were supported by NIH grant HL-68922. The OMG-SCD study has been supported by grant awards 2015131 and 2012126 from the Doris Duke Charitable Foundation, R01HL68959 and R01HL079915 from the National Heart, Lung and Blood Institute, and DK110104 and DK124836 from the National Institute of Diabetes and Digestive and Kidney Diseases.

## Author contributions

CB contributed to PathQuant package implementation with validation and support from RP and PM, supervised by JGH. SC and STL developed and implemented an earlier version of the PathQuant package and performed preliminary analyses of the TK study, supervised by MR, JDR, GL and CDR. CB performed pre-processing of all mGWAS result datasets, generated final results using PathQuant, interpreted the results and generated all figures and tables, supervised by CDR, MR and JGH. GL, MJT, PB and AEAK generated the data from the SCD study. YB and MEG performed the mGWAS of the SCD cohort, supervised by GL. CB and JGH wrote the manuscript, with major input from MR and CDR. All authors revised and approved the final version of the manuscript.

## SUPPLEMENTAL INFORMATION

### Supplementary figures and legends

**Supplementary Figure 1.**
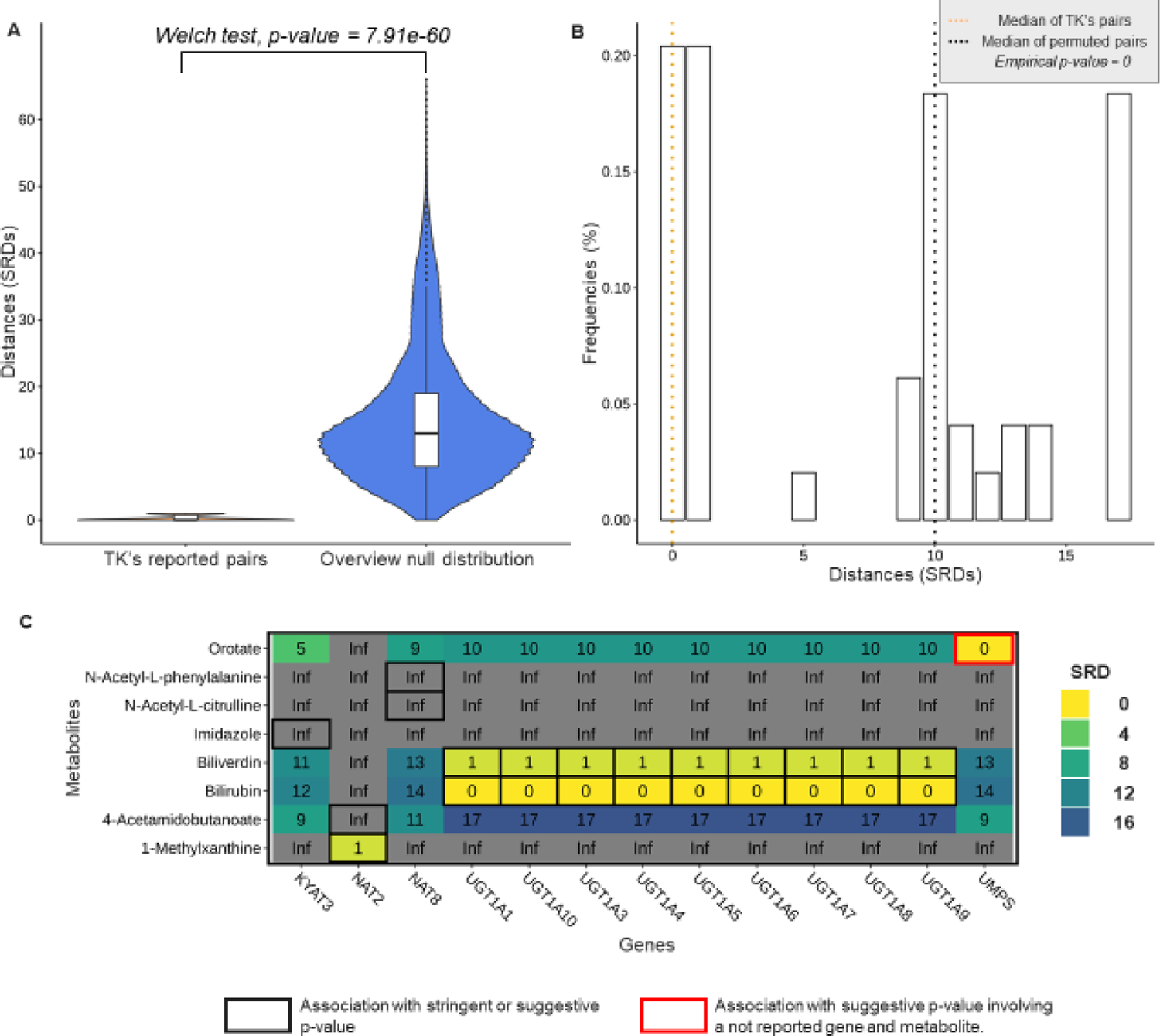
SRD of stringently and suggestive associated gene-metabolite pairs in the HMC study. (A) Comparison of SRD annotations for reported genome-wide significant associations from the HMC study (orange) and the distribution of all SRD values within KEGG overview graph (hsa01100) (B) Distribution of SRD values computed from permuted gene-metabolite pairs from HMC study, with a median SRD of 10 (black dotted line). The median SRD of 0 (orange dotted line) represents an empirical p-value of p=0. (C) Heatmap representing all genes and metabolites included in reported stringent and suggestive associations from the HMC study. The 24 mapped gene-metabolite associations are enclosed: a black box indicates a reported stringent or a suggestive pair for which either the gene, the metabolite or both have been reported in the dataset (n=23) while a red enclosed box indicates a suggestive pair for which the gene and the metabolite have not been reported in the study (n=1).

**Supplementary Figure 2.**
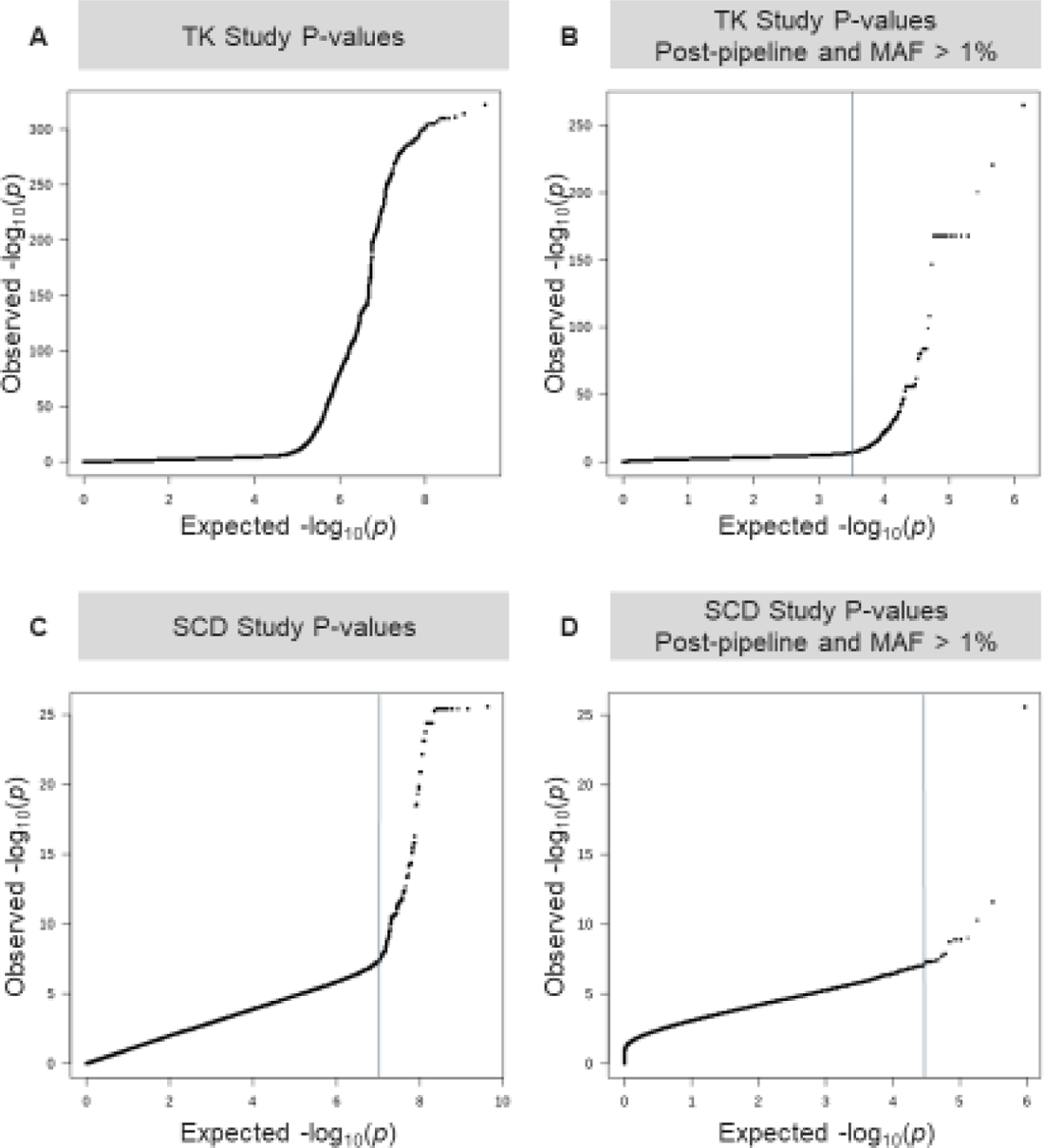
QQplots for the TK and SCD studies. (A/C) QQplots using all p-values from mGWAS files available, before any pre-processing steps for TK study (A) and for the SCD study (C). (B/D) QQplots using p-values at the end of the pipeline (See Methods) for the TK study (B) and for the SCD study (D). Vertical lines indicate the graphically determined cut-offs we used in our Results section. Abbreviations: MAF = Minor Allele Frequency, QQplot = Quantile-Quantile plot.

**Supplementary Figure 3.**
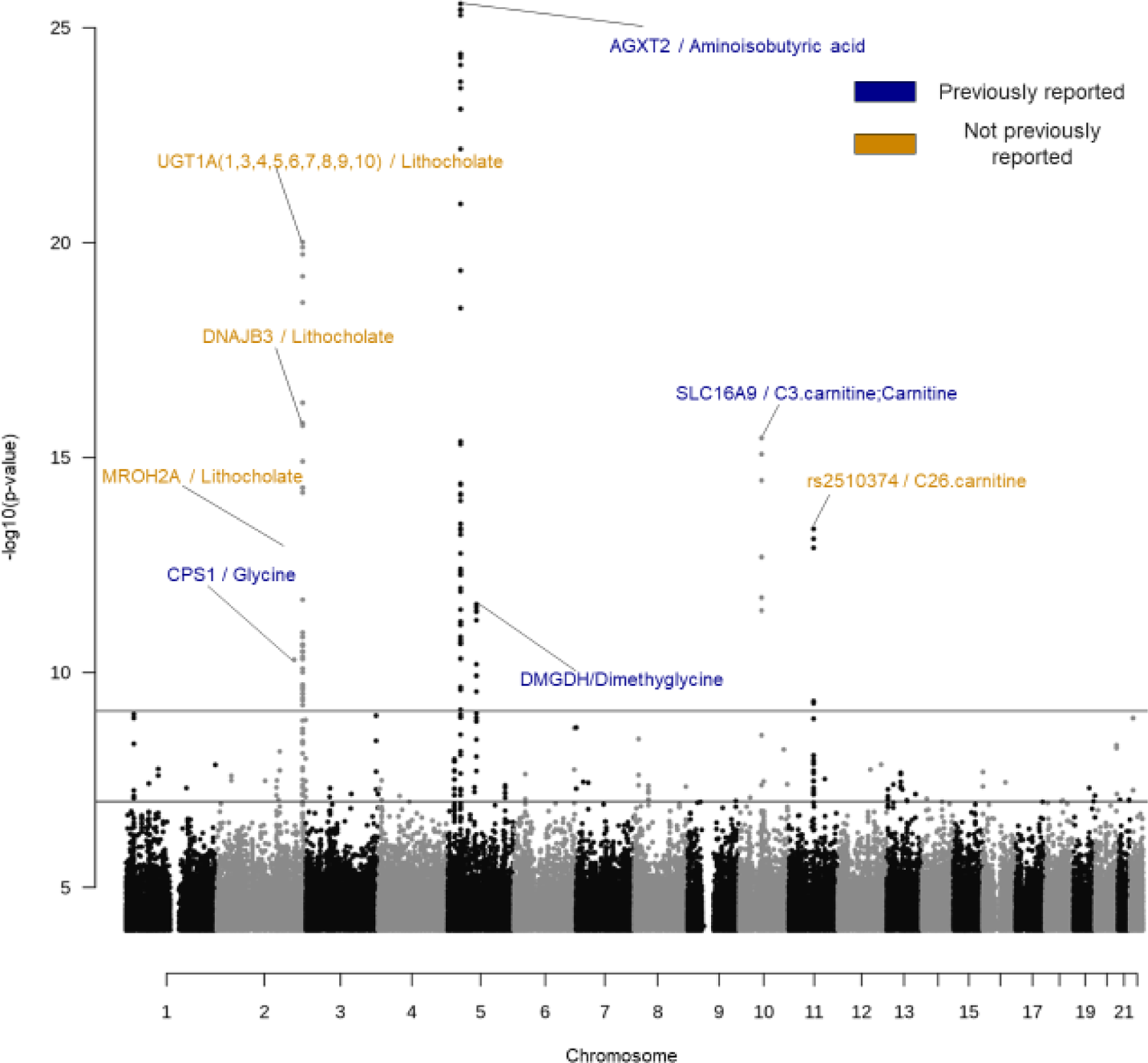
Manhattan plot of the SCD study. Manhattan plot of the SCD study with a p-values below 1×10^-4^ cutoff, each dot represents an association between a variant and a metabolite. Black and white is an alternating color palette for chromosomes. The first line from plot bottom (dark plain) indicates the genome-wide threshold at 1×10^-7^, and the second line (light plain) at 7.8125×10-10, indicates the first threshold corrected by Bonferroni (See Methods). Top hits for each mapped gene above the Bonferroni threshold are labelled with two distinct colors: blue if the association has not been previously reported, or orange if the association has been already reported in the literature. Gene symbols are separated to metabolite name(s) by “/” symbol.

**Supplementary Figure 4.**
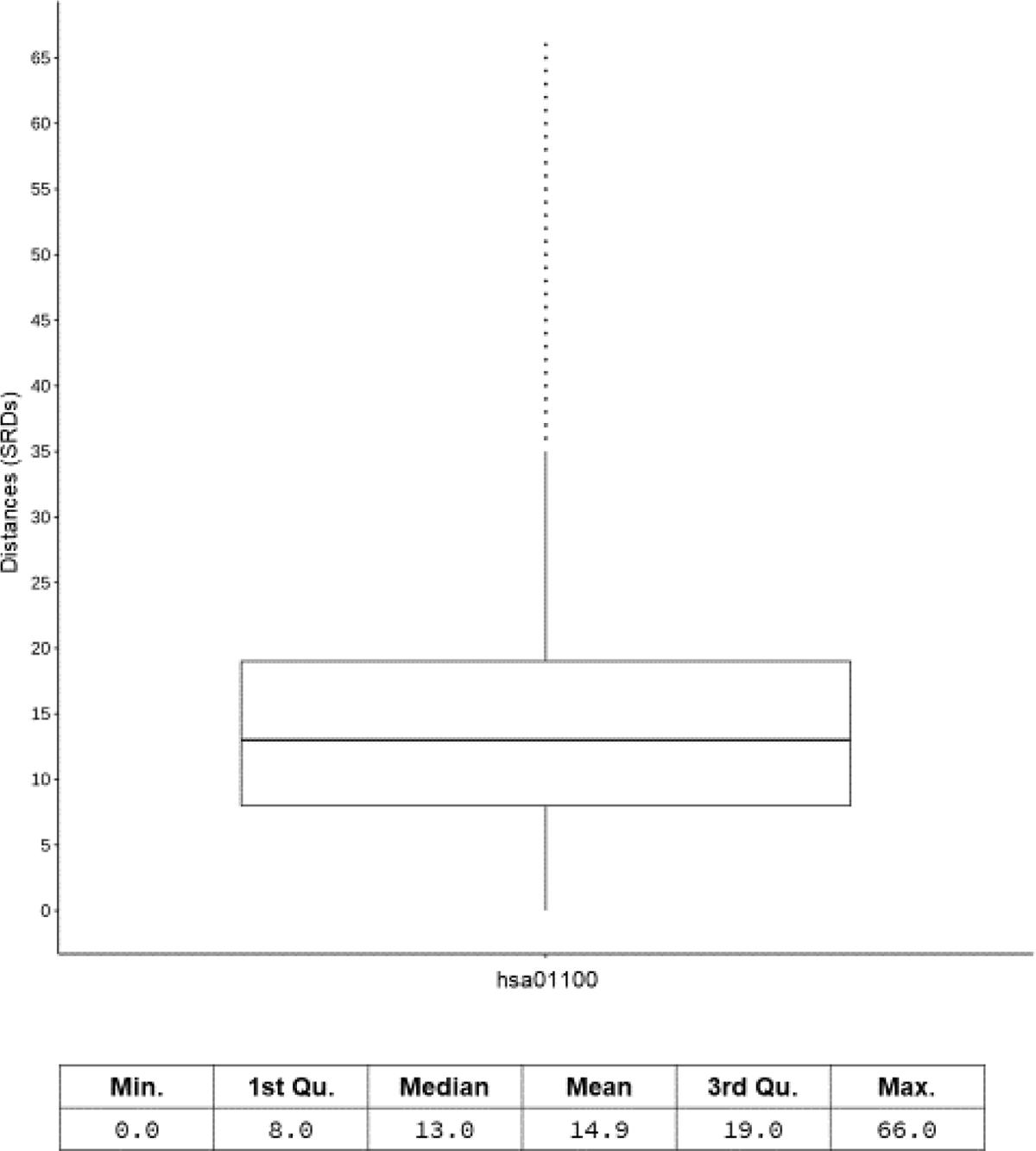
Summary statistics of the null distribution of SRD values within the KEGG overview graph. Boxplot of the SRD values and table of summary statistics for the null distribution of SRD values within the KEGG overview graph (hsa01100). The mean SRD within this graph is 14.89, with a maximum value of 66. The first quartile (SRD <8) is used as the threshold to categorize any pair with a close or far biological relationship label.

## SUPPLEMENTARY METHODS

### PathQuant tool to compute shortest reactional distances (SRD)

The method was developed using R and delivered as an R package. PathQuant (for Pathway Quantity).

### 1.1 Input parameters

#### Gene-metabolite pairs

PathQuant accepts the following input for SRD computation: 1) gene(s); and 2) their associated metabolite(s) in columns, each with their specific KEGG identifiers (IDs). Each row represents a unique gene-metabolite association. Only associations between genes and individual metabolites are taken as entry, hence associations between a gene and a ratio of metabolites must be separated prior to analysis (usually by generating gene-numerator/denominator pairs).

#### Selected metabolic pathways

PathQuant accepts a list of metabolic pathways, each with its specific KEGG IDs. For this application, we used the map ID: hsa: 01100 in KEGG (referred herein as the ‘overview’ or ‘hsa01100’ throughout this article).

#### 1.2 Association classification

PathQuant can classify each association of the input by gene product into four broad categories: enzyme, transporter, other (other proteins, transcription factor, and more) and not classified using KEGG Brite database.

#### 1.3 Metabolic network modelling

The selected KEGG pathway, encoded in KEGG XML file format (KGML), is downloaded using the KEGG API, from the most up-to-date KEGG pathways available, and then moved to a specific version folder in order to keep track, or use a specific version of interest. Users can choose which downloaded version to use or let PathQuant use the up-to-date version. The pathway is then converted into a graph of biochemical reactions (also called compound graph) with metabolites, as nodes and genes, mapped to their corresponding encoded enzymes, as edges. The topology of the pathway is captured in the constructed graph. Genes encoding enzymes catalysing multiple reactions are mapped to multiple edges. The constructed graph represents exactly the metabolic pathways of KEGG which are built mainly with metabolites that are the main reactants of a reaction, dismissing cofactor metabolites, such as NAD, or common co-substrates/products, such as ATP or H2O. Finally, we use a non-oriented graph as the KEGG standards are not consistent in this matter.

#### 1.4 SRD computation

Our method computes the SRD, which is defined as the shortest reactional distance path between a given gene and a metabolite. The SRD is computed using each metabolic pathway received in input, in which a given pair is mapped. The SRD is computed as described in the workflow figure (Figure1): A distance of 0 is assigned to metabolites, which are the main substrates or products of the reaction catalysed by the enzyme encoded by the selected gene of interest. The SRDs to all other metabolites are obtained using the breadth-first search algorithm. The algorithm is used to find the SRD starting from the substrate and from the product of the mapped gene to the paired metabolite, thereby selecting the smallest SRD between these two. Figure1 depicts an example of SRD computation for a hypothetical reaction The main reactants of this reaction: the substrate and the product are set at an SRD of 0. Everything running deeper than the substrate and the product within the graph is adding a distance of one for each depth: SRD = 1 for the green hypothetical metabolites and SRD = 2 for the blue hypothetical metabolite.

#### 1.5 SRD metric analysis

The utility of the computed SRD metric for the annotation of gene-metabolite associations reported by mGWAS was assessed using different approaches (See Methods).

#### 1.6 Data outputs and visualisation

PathQuant outputs a text file containing gene and metabolite classification, Enzyme Commission number (EC), KEGG Brite, KEGG IDs of used pathways for the SRD computation, and SRD values for all associations. These SRD values can also be visualised in a heatmap and global or multiple distribution plots; a few examples are available at in this manuscript (Results).

### TK datasets and naming

For the TK study, we downloaded the file named “NIHMS58114-supplement-2.xlsx” from the supplementary section of the publication. This first file contains only stringently associated pairs. We make a distinction between the TK and the “TK None” datasets as they do not use the same files from the original publication. The TK None is referring to the GWAS summary stats, without any p-value cut-off, directly downloaded from here: https://metabolomics.helmholtz-muenchen.de/gwas/index.php?task=download. We downloaded and merge the files named: shin_et_al.metal.out.tar.gz, shin_et_al.xeno.metal.out.tar.gz. As the positions were coming from NCBI Build 36, we performed a lift over to hg19 in order to have same build across all different studies.

