## Supplementary material for "Gene-metabolite annotation with shortest reactional distance enhances metabolite genome-wide association studies results": Key Resources Table

| REAGENT or RESOURCE | SOURCE | IDENTIFIER |
|---|---|---|
| **Deposited Data** | | |
| TK | https://doi.org/10.1038/ng.2982 | https://static-content.springer.com/esm/art%3A10.1038%2Fng.2982/MediaObjects/41588_2014_BFng2982_MOESM50_ESM.xlsx |
| TK None | https://doi.org/10.1038/ng.2982 | Summary stats without p-value cutoff: https://metabolomics.helmholtz-muenchen.de/gwas/ |
| HMC | https://doi.org/10.1038/s41467-017-01972-9 | Supplementary 7, https://static-content.springer.com/esm/art%3A10.1038%2Fs41467-017-01972-9/MediaObjects/41467_2017_1972_MOESM9_ESM.xlsx |
| SCD | 10.3324/haematol.2022.281180 and 10.1016/j.bcmd.2020.102504 | https://github.com/HussinLab/PathQuant/blob/main/Publication/SCD_study_download_link.txt |
| Supplementary data | To be submitted | Available at www.github.com/HussinLab/PathQuant/Publication/References DOI on Zenodo will be provided upon acceptance. |
| **Software and Algorithms** | | |
| PathQuant | This study | www.github.com/HussinLab/PathQuant/ |
| bedtools version v2.30.0 | https://doi.org/10.1093/bioinformatics/btq033 | www.github.com/arq5x/bedtools2/releases/tag/v2.30.0 |
| R statistical software 3.4.4 | R Foundation for Statistical Computing | www.r-project.org/ |
